## Supplementary figures and images for "Circular RNA *HMGCS1* sponges *MIR4521* to aggravate type 2 diabetes-induced vascular endothelial dysfunction"

### Figure 1-figure supplement 1

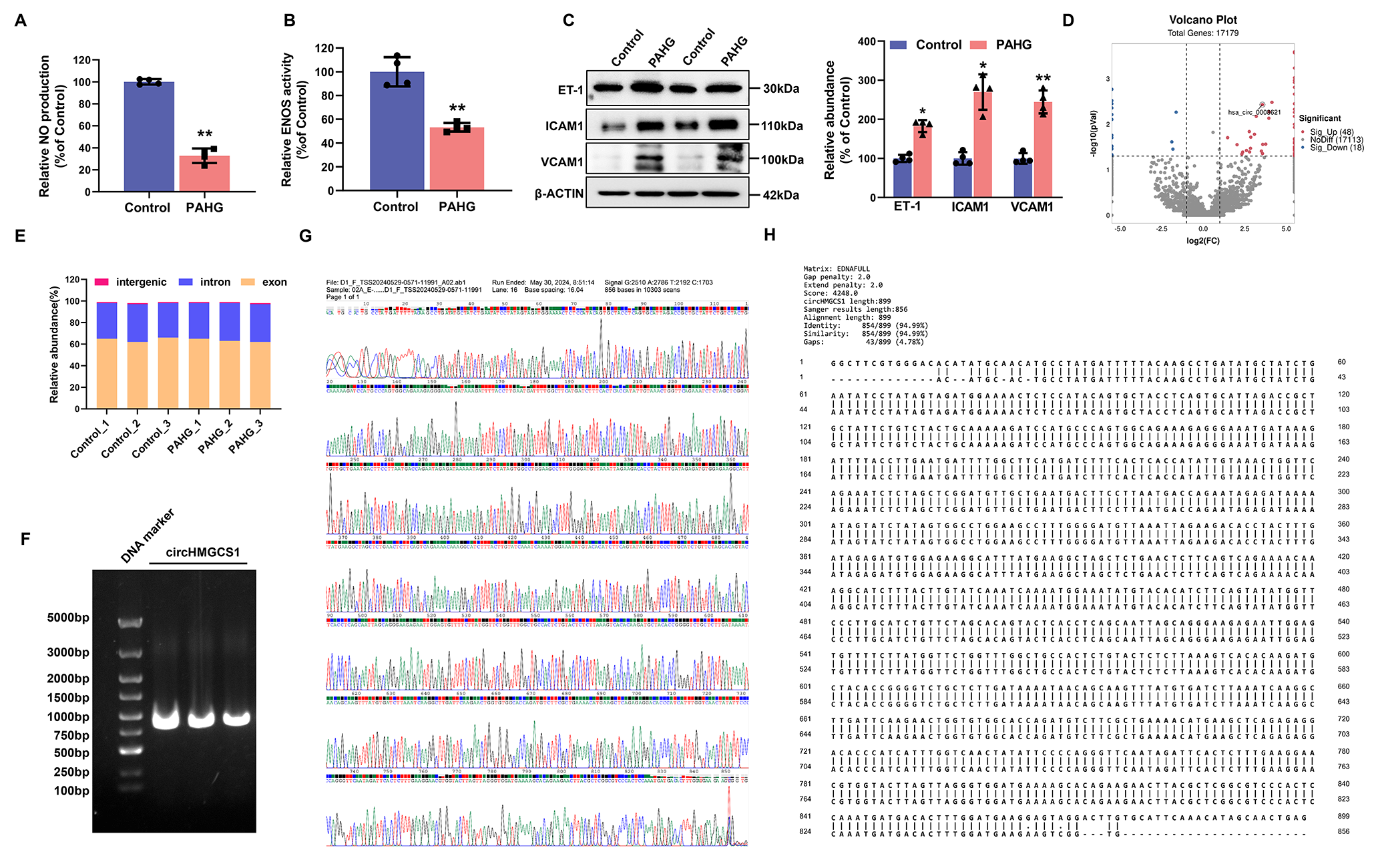

### Figure 2-figure supplement 1

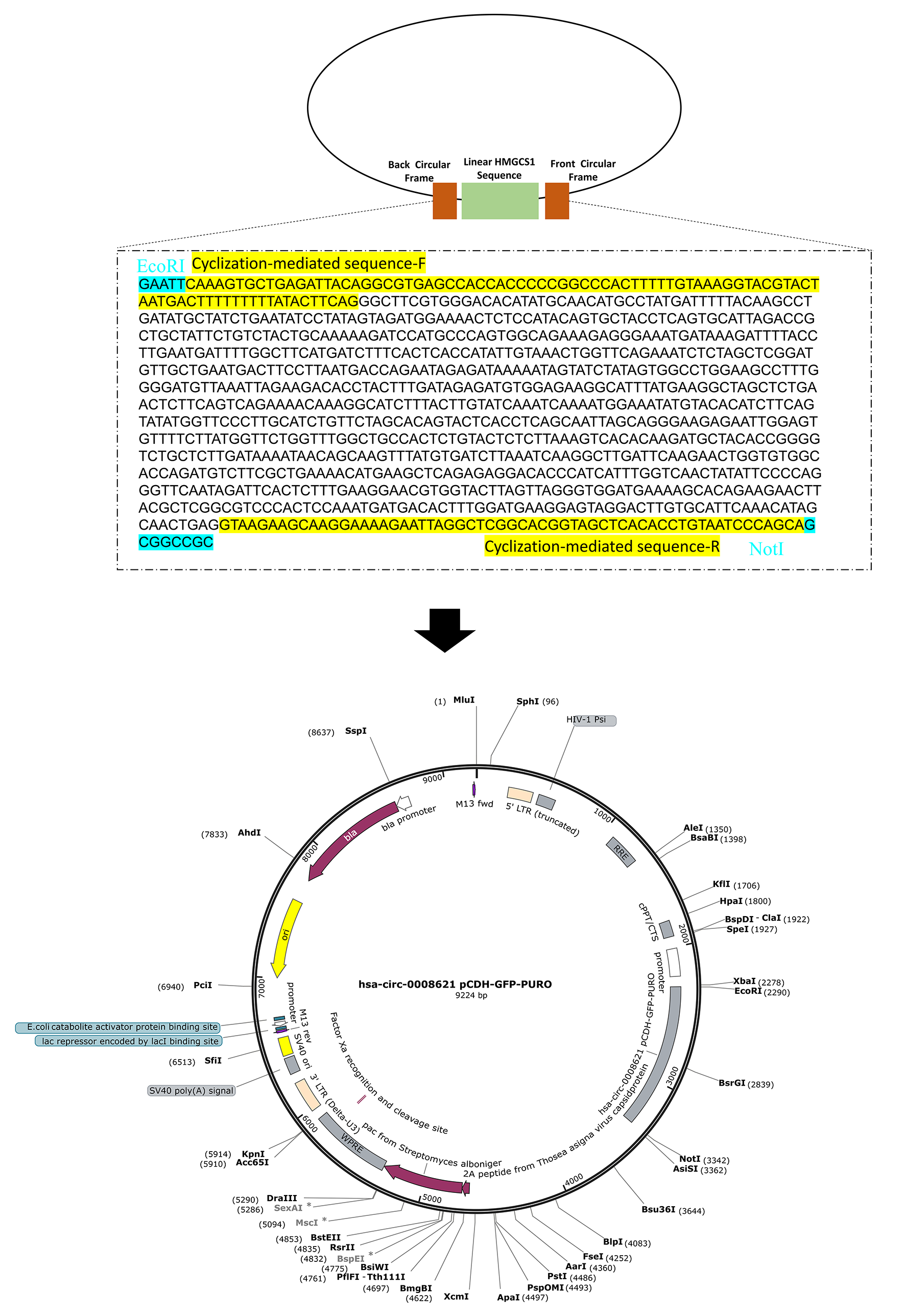

### Figure 3-figure supplement 1

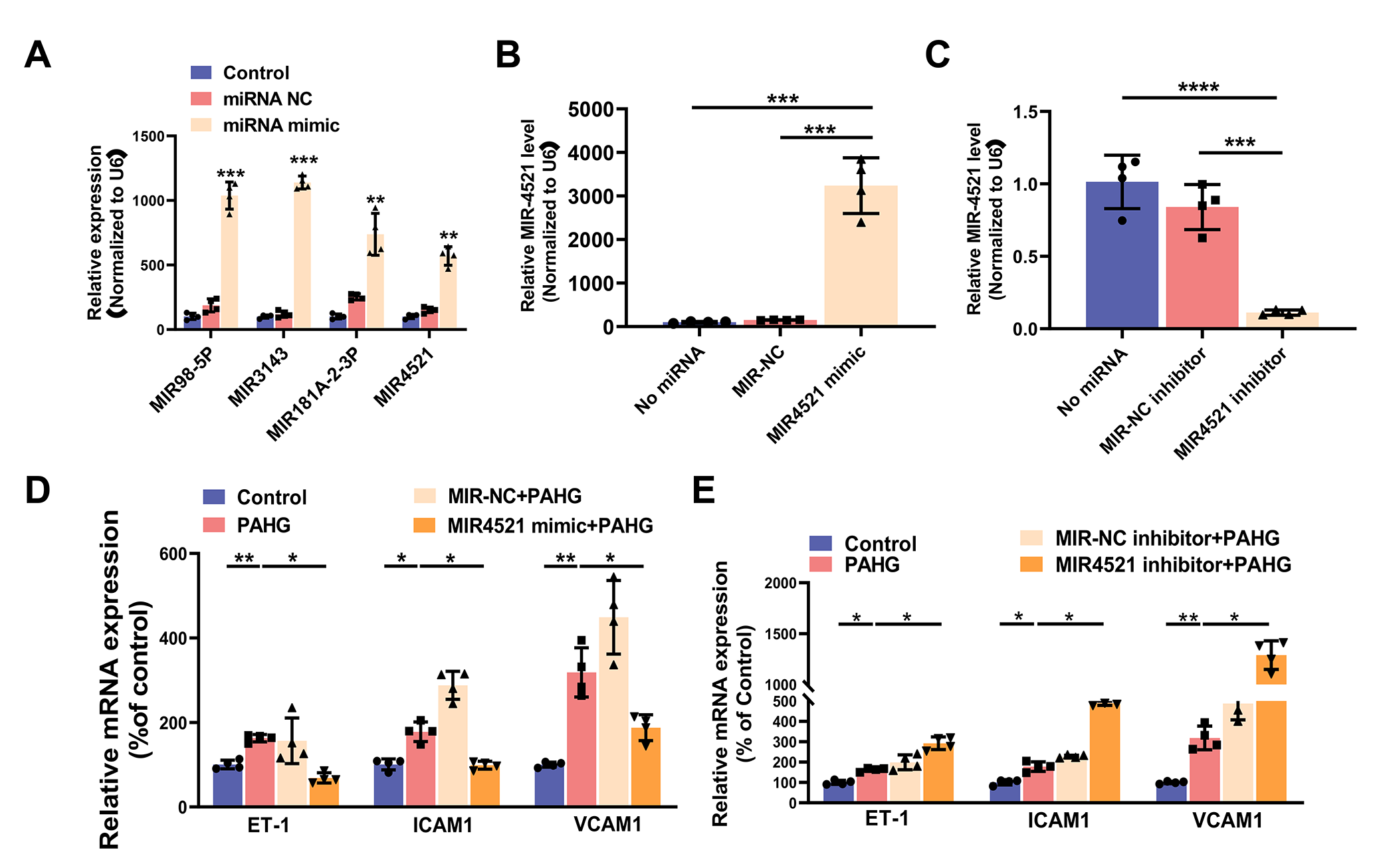

### Figure 4-figure supplement 1

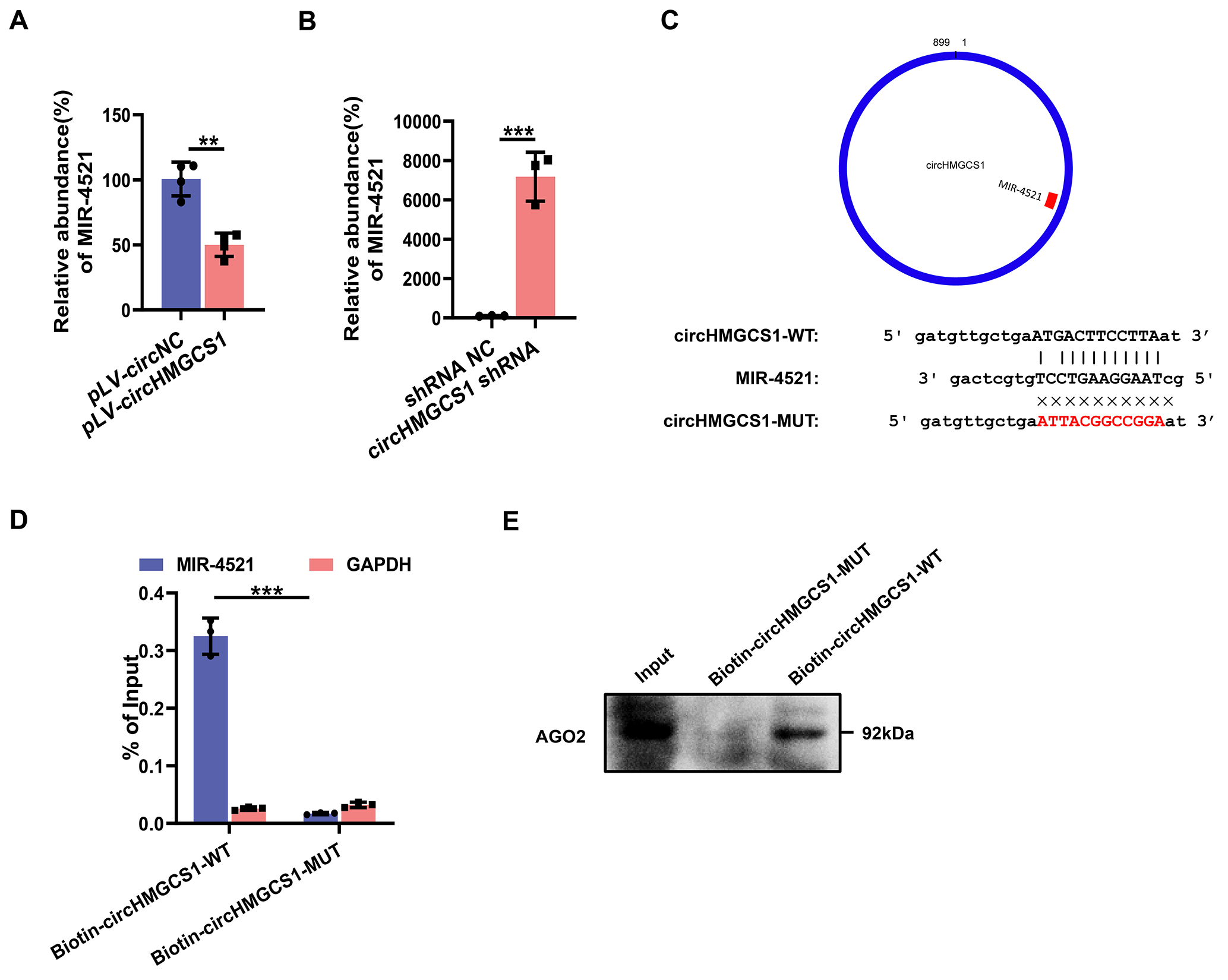

### Figure 5-figure supplement 1

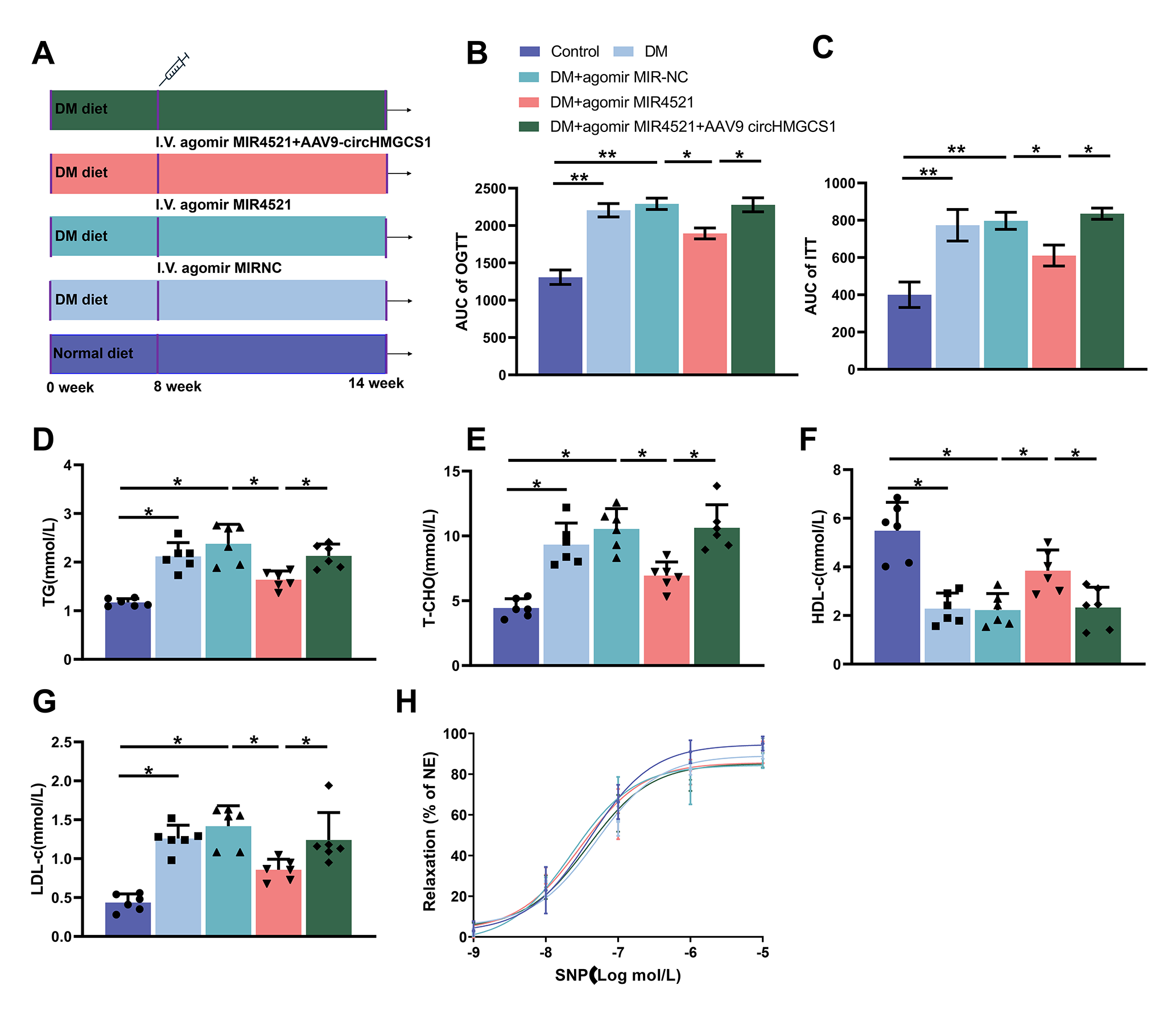

### Figure 6-figure supplement 1

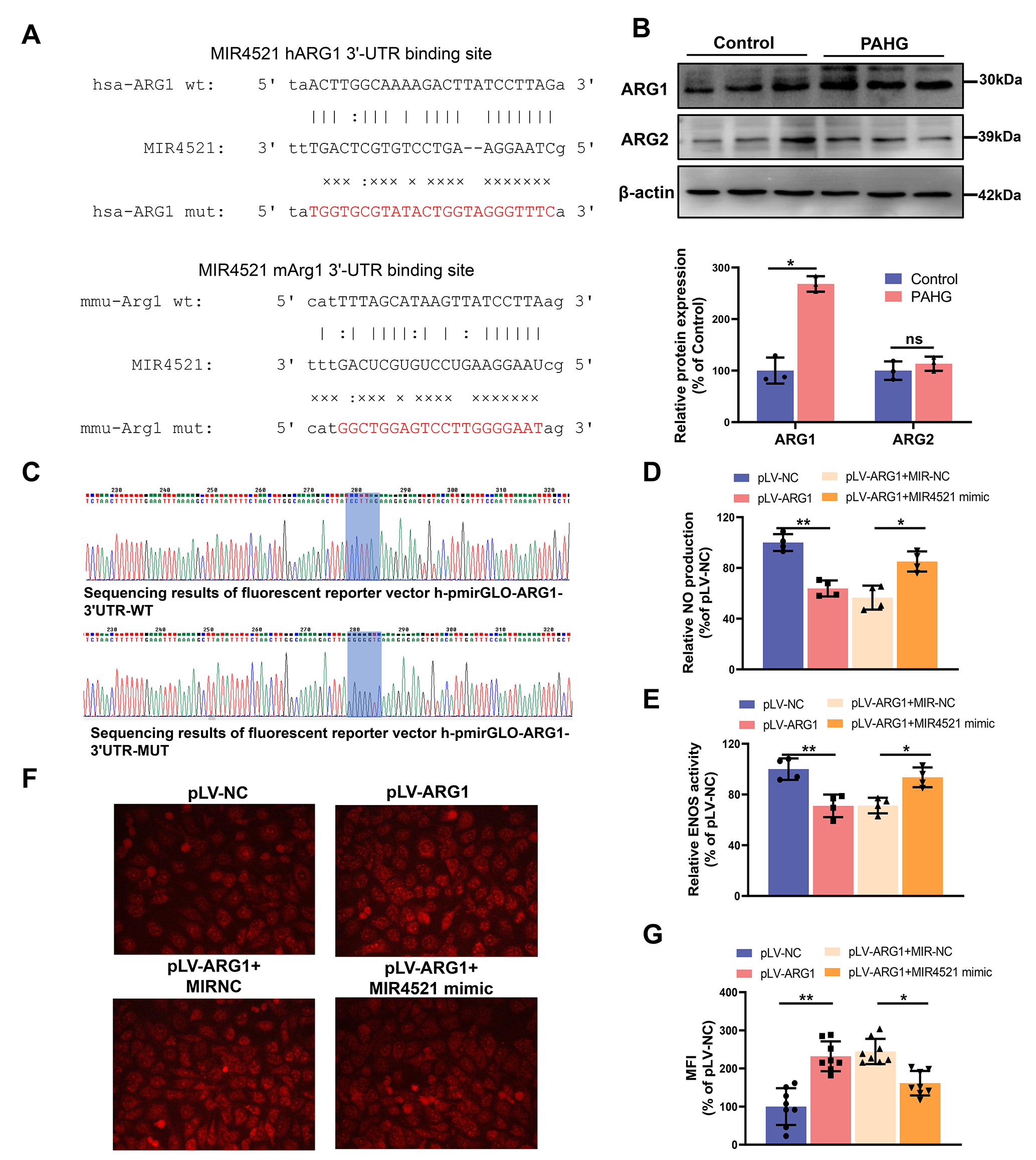

### Figure 7-figure supplement 1

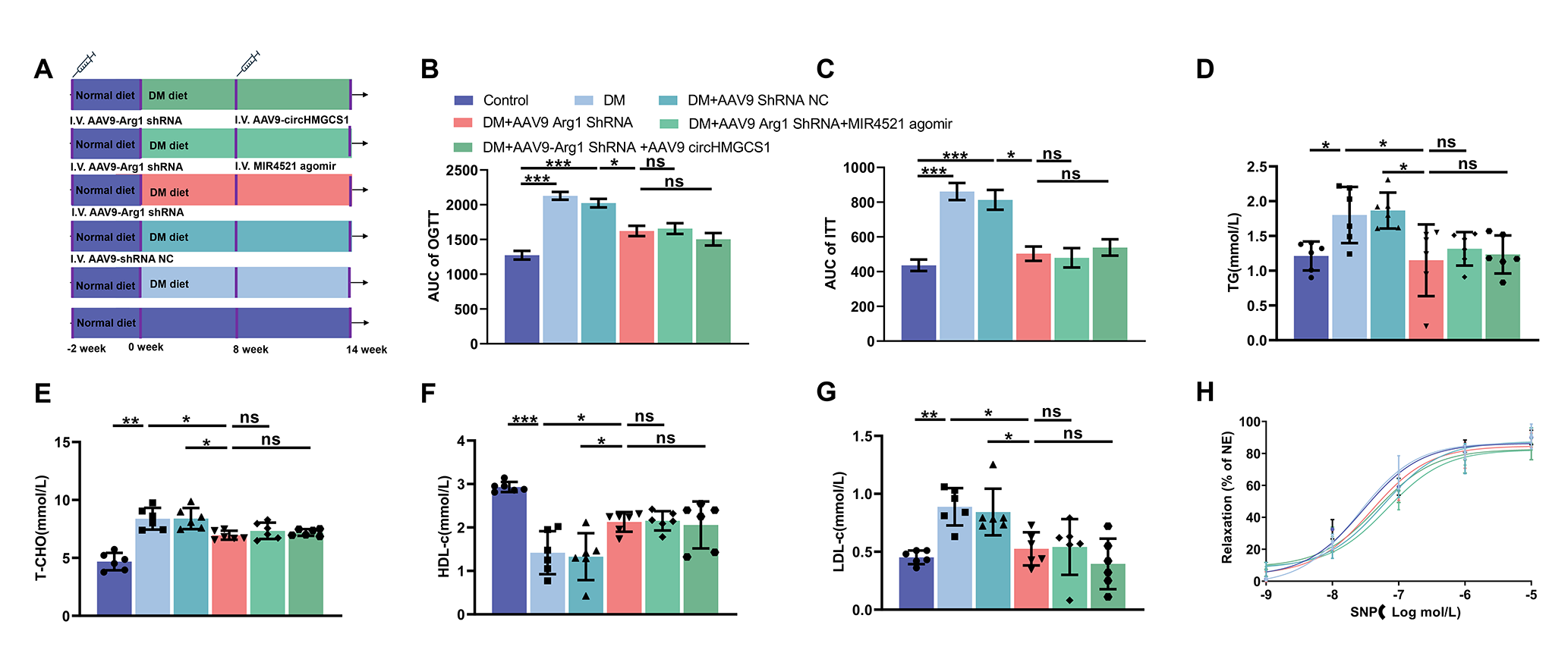
